# Coral venom and toxins as protection against crown-of-thorns sea star attack

**DOI:** 10.1101/2025.02.10.637440

**Authors:** Lucy M. Gorman, Ariana S. Huffmyer, Maria Byrne, Suzanne C. Mills, Hollie M. Putnam

**Affiliations:** PSL Université Paris: EPHE-UPVD-CNRS, USR 3278 CRIOBE, BP 1013, 98729 Papetoai, Moorea, French Polynesia; Laboratoire d’Excellence “CORAIL”, France; Biological Sciences, University of Rhode Island, Kingston, RI 02881; School of Aquatic and Fisheries Sciences, University of Washington, Seattle, WA 98195; School of Life and Environmental Sciences, Marine Science Institute, The University of Sydney, New South Wales, Australia

**Author notes:** Corresponding author: Lucy Gorman.

**Keywords:** crown-of-thorns sea star, venom, pore-forming toxins, *Acropora*, *Porites*, neurotoxins

## Abstract

Crown-of-thorns sea star (CoTS) outbreaks are one of the leading causes of hard coral cover decline across the Indo-Pacific, posing a major threat to the health and resilience of coral reefs. However, the drivers underlying feeding on preferred (e.g., *Acropora* spp.) versus non-preferred (e.g., *Porites* spp.) are poorly understood. We hypothesised that coral venom may influence CoTS food preferences. We investigated whether coral venom toxin and peptide families may drive CoTS prey preferences by comparing the genomes and transcriptomes of preferred (five *Acropora* species) and non-preferred (five *Porites* species and *Echinopora lamellosa*) prey species of CoTS. We constructed databases of known cnidarian venom toxins, and along with the full UniProtKB/Swiss-Prot Tox-Prot database, used these to identify toxin peptides and investigate function and phylogeny. The most abundant toxins across all coral species included kunitz-type neurotoxins, neurotoxic turripeptides, snake venom lectins, toxic proteases and actinoporins. There were proteins present only in certain *Porites* species but completely absent from all *Acropora* species (e.g., tereporin/conoporin, snake venom peptides) and *vice versa* (e.g., sarafotoxin). Further, *Porites* species contained a homolog to conkunitzin, a toxin known to disintegrate the tube feet of CoTS, suggesting a potential mechanism for their lower susceptibility to predation. We also observed a greater diversity of jellyfish-like proteins in CoTS-exposed *Porites* species compared to naïve *Porites* species, suggesting these proteins deter CoTS. These findings have direct applications to assessing reef coral’s susceptibility to future CoTS outbreaks and active reef management.

## 1. Introduction

Crown-of-thorns sea stars (CoTS) are found throughout the Indian and Pacific Oceans, as well as the Red Sea and the Gulf of Oman (Pratchett *et al*., 2014). Throughout these locations, four to five CoTS species have been identified, which comprise the *Acanthaster* species complex: including the Northern Indian Ocean species *Acanthaster planci sensu strictu*; the Red Sea species *Acanthaster benziei;* and the Southern Indian Ocean species *Acanthaster mauritiensis* (Vogler *et al*., 2008; Haszprunar *et al*., 2017; Wörheide *et al*., 2022; Foo *et al*., 2024; Uthicke *et al*., 2024). The taxonomy of the Pacific species is uncertain with *Acanthaster* cf. *solaris* as a potential name for the western Pacific species and *Acanthaster* cf. *ellisii* in the eastern Pacific (Uthicke *et al*., 2024). These sea stars are corallivorous (review Foo *et al*., 2024). At low densities (1-10 sea stars *per* hectare; Pratchett *et al*., 2014; Dumas *et al*., 2016), CoTS are argued to be beneficial to reefs, as they preferentially prey on faster-growing coral species (e.g. *Acropora* species), creating more space for slower-growing species (e.g. *Porites* species), and thus, increase the coral diversity (Done & Potts, 1992; Bellwood *et al*., 2024). In contrast, high density “outbreak” populations of CoTS where densities can reach up to 15,000 sea stars *per* hectare can decimate reefs (Dumas *et al*., 2022). Each CoTS is able to consume 5-12 m^2^ of coral surface annually (Chesher, 1969; Pearson & Endean, 1969; Dana & Wolfson, 1970), and outbreak populations spread at speeds of up to 60 km *per* year (Reichelt *et al*., 1990; Vanhatalo *et al*., 2017). These outbreaks drive significant declines in coral cover (Osborne *et al*., 2011; De’Ath *et al*., 2012), compounding the impacts of heatwave-induced bleaching, which has now become the primary driver of coral mortality (Bozec *et al*., 2022; Byrne *et al*., 2025).

Early post settlement, CoTS juveniles are herbivores with a preference for crustose coralline algae (CCA) and under favourable conditions, they undergo an ontogenetic diet transition to coral prey when they reach around 8 mm diameter (Yamaguchi, 1974). After this ontogenetic diet transition, their size rapidly increases (Zann *et al*., 1987; Deaker *et al*., 2020). In order to locate their coral prey, CoTS use olfactory sensing (Ormond *et al*., 1976; Ling *et al*., 2020; Webb *et al*., 2024), with the densities of preferred coral prey shown to influence the behaviour of CoTS to either vacate or stay on a reef (Ling *et al*., 2020). Where *Acropora* and pocilloporid corals (e.g. *Seriatopora* and *Stylophora* species) are available, CoTS exhibit a preference for these as prey (De Bruin, 1972; Ormond *et al*., 1973; Keesing, 1990; De’ath & Moran, 1998; Johansson *et al*., 2016; Keesing, 2021; Foo *et al*., 2024). They also grow faster on a diet of these corals (Keesing & Halford, 1992; Keesing, 2021). *Porites* species are among the least preferred prey of CoTS (De’ath & Moran, 1998; Pratchett, 2007; Kenyon & Abey, 2009; Millican *et al*., 2024) and CoTS have minimal growth rates on a diet of these corals (Keesing, 2021). Even at very low abundances of preferred coral prey (*Acropora* and pocilloporid corals), CoTS still show preference for these species, albeit with increased consumption of non-preferred coral taxa (Keesing *et al*., 2019; Keesing, 2021).

It is still unclear why CoTS show feeding preferences for certain coral taxa. Hypotheses include: nutritional differences between corals (Ormond *et al*., 1976; Keesing, 1990); accessibility to coral tissue based on coral morphotype (e.g. tabular *versus* branching) and ease of tissue digestion (Keesing, 1990); coral symbionts that repel CoTS (shrimps, gobies, trapeziid crabs) (Glynn, 1980; Pratchett, 2001; also see Montano *et al.,* 2017); and, coral nematocyst and venom defence (Barnes *et al*., 1970; Ormond *et al*., 1976; Deaker *et al*., 2021). Recent studies are starting to unveil the interplay between CoTS and coral prey choice. For instance, a laboratory study concluded that CoTS’ preference for *Acropora* species is not driven by growth form or nutritional content, but rather by prior exposure to that coral species during rearing (Johansson *et al*., 2016). This suggests that feeding preference may involve a conditioning process, potentially akin to an immune-type response. Furthermore, as CoTS mature they increase their ability to feed on a wider range of corals, including those that can cause damage to them as juveniles (Johansson *et al*., 2016). These findings support the hypothesis that CoTS feeding preferences are influenced by their ability to tolerate or adjust to coral venom defences as a result of previous exposure. This is particularly relevant given the dose-dependent nature of venom bioactivity, which may explain why larger juveniles can diversify their diet (Deaker *et al*., 2021), as venom is less likely to cause lethal injuries at this stage.

Coral venom and toxins provide protection against predation. For instance, a toxin isolated from the soft coral *Sarcophyton glaucum*, sarcophine, deters predatory corallivorous fishes (Ne’eman *et al*., 1974). These toxins may allow corals to cause lethal and sub-lethal injuries to CoTS during contact and ingestion, with the coral *Echinopora lamellosa* causing 100% mortality of CoTS post-ingestion (Johansson *et al*., 2016). Furthermore, CoTS exhibit a humped posture during feeding to protect their tube feet, which are more vulnerable to damage from coral nematocysts than the stomach (Barnes *et al*., 1970). On smaller coral colonies specifically, this humped posture allows the sensitive tube feet to remain on a non-nematocyst surface (e.g., calcareous rock), whilst exposing the less-sensitive stomach to the coral (Barnes *et al*., 1970). This theory is further supported by observations of discharged nematocysts on CoTS tube feet after exposure to coral (Moore *unpublished*, as cited in Moore & Huxley, 1976). Furthermore, venom defence from other cnidarian taxa such as hydrozoans may provide protection for the coral against CoTS e.g., neighbouring *Millepora* species have been shown to produce the strongest aversion reaction in CoTS (Moore & Huxley, 1976) and may provide refuge for *Acropora* species, against CoTS predation (Kayal & Kayal, 2017). It has also been suggested that the venom from *Acropora digitifera* has a defensive role against predators due to the low diversity and abundance of toxins present in the venom, contrastingly to predatory venoms, which have a larger suite of toxins due to high evolutionary selection pressure (Gacesa *et al*., 2015). These studies indicate that investigating the venom toxin repertoire of CoTS prey may be imperative to understanding their resilience against CoTS attack.

Cnidarian venoms are understudied compared with those of other venomous animals such as snakes and cone snails, for which the biochemistry and molecular biology of the bioactive peptides are well understood (Turk & Kem, 2009). Within Cnidaria, the toxin repertoire of jellyfish and sea anemones has been a focus point of research (Gacesa *et al*., 2015; Jouiaei *et al*., 2015; Schmidt *et al*., 2019; Klompen *et al.,* 2022), with the venom of sea anemones harbouring one of the most diverse array of toxins polypeptides found across the animal kingdom (Castañeda & Harvey, 2009; Frazão *et al*., 2012; Orts *et al*., 2013; Finol-Urdaneta *et al*., 2020; An *et al*., 2022). Studies of scleractinian venoms show that they are all rich in biologically active peptides (Ne’eman *et al*., 1974; Radwan *et al*., 2002; Gacesa *et al*., 2015; Garcia-Arredondo *et al*., 2016; Ben-Ari *et al*., 2018; Yosef *et al*., 2020; Drake *et al*., 2021; Schmidt *et al*., 2022), with significant similarities in the major classes of toxins present across Cnidaria (Rachamim *et al*., 2015; Gacesa *et al*., 2015). For instance, some commonly found toxins and peptide/protein families in cnidarians include: cytolysins, including but not exclusive to, pore-forming toxins (PFTs; e.g., aerolysins, actinoporins and Membrane Attack Complex/ Perforin toxins (MAC-PF)), neurotoxins (e.g., K_V_/Na_V_ channel neurotoxins and the peptidic neurotoxic small cysteine rich proteins (SCRiPS)), and cytotoxic enzymes (e.g., peptidases, metalloproteases and phospholipase A2s (PLA2)). The most abundant toxins across cnidaria and their known function are described below in **Table 1**.

**Table 1.** Most common toxins in cnidarian venoms and their cellular functions/bioactivity.

| Toxin | Function/Bioactivity |
| --- | --- |
| Actinoporins | A class of pore forming toxins (PFTs) which bind specifically to sphingomyelin lipids in the cell membrane and initiate the formation of a pore, leading to osmotic imbalance and ultimately, necrosis (Frazão *et al.*, 2012; Podobnik & Anderluh, 2017). Actinoporins are hypothesised to comprise most of the lethal bioactivity in cnidarian venoms (Ben-Ari *et al.*, 2018). Actinoporins contain a conserved domain known as the sea anemone cytotox domain which harbours a N-terminal amphiphilic α-helix region that is essential for pore forming activity (Anderluh *et al.*, 1997). Actinoporins also usually contain a conserved RGD-motif Arg-Gly-Asp that promotes attachment to integrin receptors on the surface of cells (Uechi *et al.*, 2010). |
| Hydralysins | Hydralysins are non-nematocyst, cytolytic and paralytic β-pore-forming toxins from green hydra *Chlorohydra viridissima* (Zhang *et al.*, 2003; Sher *et al.*, 2005a, b and are similar in activity and structure to bacterial and fungal toxins (Sher *et al.*, 2005a, b). |
| Membrane-attack complex/perforins (MACPFs) | MAC-PFs are the largest superfamily of pore-forming toxins with roles in the immune system of mammals in targeting pathogens, while mammalian parasites use these proteins to invade and infiltrate host tissues (Kondos *et al.*, 2010; Lukyanova *et al.*, 2015). Function of MAC-PF toxins in animal venoms is similar to that of MAC-PFs in the host immune system, forming pores in target membranes (victims of envenomation), leading to cell lysis and death (Sato *et al.*, 2007; Frazão *et al.*, 2012). See Nagai *et al.*, (2002); Oshiro *et al.*, (2004); Rachamim *et al.* (2015); Drake *et al.*, (2021) for studies discovering MAC-PFs. |
| *Chironex fleckeri*-like toxins (CFXs) | Jellyfish pore-forming toxins and are proposed to be the main reason behind toxicity following human envenomation (Jouiaei *et al.*, 2015). Envenomation from these toxins can lead to hemolyticity, cardiotoxicity, cutaneous inflammation, and necrosis (Jouiaei *et al.*, 2015). Also found in other cnidarian venoms apart from jellyfish e.g., ceranthids (Klompén *et al.*, 2020). |
| Kunitz-like domain neurotoxins | There are two types of Kunitz-type proteins: potent protease inhibitors and potassium channel blockers (Bayrhuber *et al.*, 2005). All kunitz-type proteins contain three highly-conserved disulfide bonds (Bayrhuber *et al.*, 2005) and can comprise some of the most lethal bioactivity in a venom e.g., dendrotoxins from black mamba venom (Laustsen *et al.*, 2015). |
| Small cysteine rich proteins (SCRiPS) | SCRiPS are neurotoxic peptides from stony corals (scleractinians) that have an eight cysteine framework, similar to the myotoxin domain of rattlesnake crotoamine toxins (Sunagawa *et al.*, 2009; Jouiaei *et al.*, 2015). SCRiPS cause significant neurotoxicity in zebrafish larvae (Jouiaei *et al.*, 2015) and have also been found in other cnidarians e.g., anthozoans (Barroso *et al.*, |
|  | 2024). |
| Shk-domain neurotoxins | Shk is a type of potent neurotoxin that targets potassium channels and was isolated from the sea anemone *Stichodactyla helianthus* (Castañeda *et al.*, 1995; Castañeda & Harvey, 2009). Shk-domain containing proteins found throughout organisms from plants to fish (Sanches *et al.*, 2024). |
| Turriptides containing the Kazal domain of protease inhibitors | Turriptides are Ion channel blockers (Hernández-Sámano *et al.*, 2020), that were originally found in turrid marine gastropods and phylogenetically similar to conotoxins in cone snails (Fassio *et al.*, 2019). Turriptides contain a conserved six cysteine residue domain (Aguilar *et al.*, 2009) and are also found in sea anemones (Mazzi Esquinca *et al.*, 2023), hydrozoa (Hernández-Elizárraga *et al.*, 2023) and jellyfish (Brinkman *et al.*, 2015; Edirisinghe *et al.*, 2024) venoms. |
| Phospholipase A2 (PLA2) | PLA2s are enzymes that catalyse the hydrolysis of lipids by cleaving glycerophospholipids at the *sn*-2 position into free fatty acids and lysophospholipids (Talvinen & Nevalainen, 2002; Nevalainen *et al.*, 2004). They can act as presynaptic neurotoxins that block nerve terminals by binding to nerve cell membranes where they hydrolyse membrane lipids, and inhibit neurotransmission (Frazão *et al.*, 2012). PLA2s are classed as a venom allergen, eliciting degranulation of mast cells and basophils in victims of insect envenomation (King *et al.*, 1978; Guido-Patiño & Plisson, 2022). PLA2s lead to systemic envenomation following contact (Nevalainen *et al.*, 2004) and can have several biological effects e.g., neurotoxic, cytotoxic and haemotoxic (Bittenbinder *et al.*, 2024). Along with metalloprotease activity, PLA2 activity is the main bioactivity in the jellyfish venom *Nemopilema nomurai* (Yu *et al.*, 2021). |
| Hyaluronidase | Hyaluronidase is a venom allergen enzyme, which like PLA2s, is widely present throughout all animal venoms and an integral venom spreading factor by degrading hyaluronan, the main glycan of the extracellular matrix in animals (Müller *et al.*, 2017).. |
| Acetylcholinesterase | Acetylcholinesterase is a neurotoxic enzyme that hydrolyses the neurotransmitter acetylcholine and in venoms leads to transmission disruption in the central nervous system and neuromuscular junction of their prey, leading to paralysis (Ahmed *et al.*, 2009). |
| Metalloproteases | In cnidarian venoms, metalloproteinases are shown to be key bioactive enzymes eliciting a myriad of envenomation toxicities (Hwang *et al.*, 2022). There are several families of metalloproteases present in venoms, e.g. astacin-like metalloproteases, zinc metalloproteinases, neprilysin-like toxins, glutamyl-peptide cyclotransferases and snake venom metalloproteinases (SVMP) (Weston *et al.*, 2013; Klompen *et al.*, 2020). Snake venom metalloproteinases (SVMPs) represent a major component of both snake and cnidarian venoms (Markland & Swenson, 2013; Weston *et al.*, 2013). In snake venoms, they are key enzymes contributing to the toxic activity of the venom and can possess hemorrhagic activity, fibrin(ogen)olytic activity, apoptotic activity (Markland & Swenson, 2013). They can |
|  | also act as prothrombin activators, activate blood coagulation factor X, inhibit platelet aggregation, are pro-inflammatory and inactivate blood serine proteinase inhibitors (Markland & Swenson, 2013; Olaoba *et al.*, 2020). |

The study of toxins and venomics is still in its infancy, with an estimated ∼29,000 undiscovered toxins in Cnidaria (Weston *et al*., 2013). In addition to knowledge gaps surrounding the specific toxins and peptides present in scleractinian venoms, the biological roles of these venoms are still unclear. Therefore, in this study we used genomic and transcriptome databases to investigate the abundance and diversity of toxins in the genomes of five species of CoTS preferred prey (*Acropora* species) and six species of non-preferred prey (*Porites* species and *E. lamellosa*). Our aim was to determine if there are differences in the toxin repertoire of CoTS preferred and non-preferred coral prey species to clarify the natural defenses that corals employ to deter CoTS and if these subsequently dictate the feeding preferences observed by CoTS. We could thus leverage this knowledge to inform the design of effective management measures to combat CoTS outbreaks (e.g., planting more venomous corals across at risk reefs), as in contrast to climate warming, CoTS outbreaks are relatively amenable to control, sparking considerable interest in developing intervention tools to suppress this species (Babcock *et al*., 2020; Harris *et al*., 2025). As venom toxins are known to be highly conserved (Weston *et al*., 2013; Brinkman *et al*., 2015; Gacesa *et al*., 2015), we used known cnidarian toxin and venom peptide families from related and more distantly related cnidarians as queries against our 11 scleractinian species. Additionally, we also used a discovery pipeline by BLASTing the UniProtKB/Swiss-Prot Tox-Prot database (a database of toxins and venom proteins from all venomous species; Jungo *et al*., 2012) to search for new putative toxin and venom peptides.

## 2. Methods

Cnidarian genome and transcriptome databases were acquired from publicly available sources (**Table 2**).

**Table 2.**
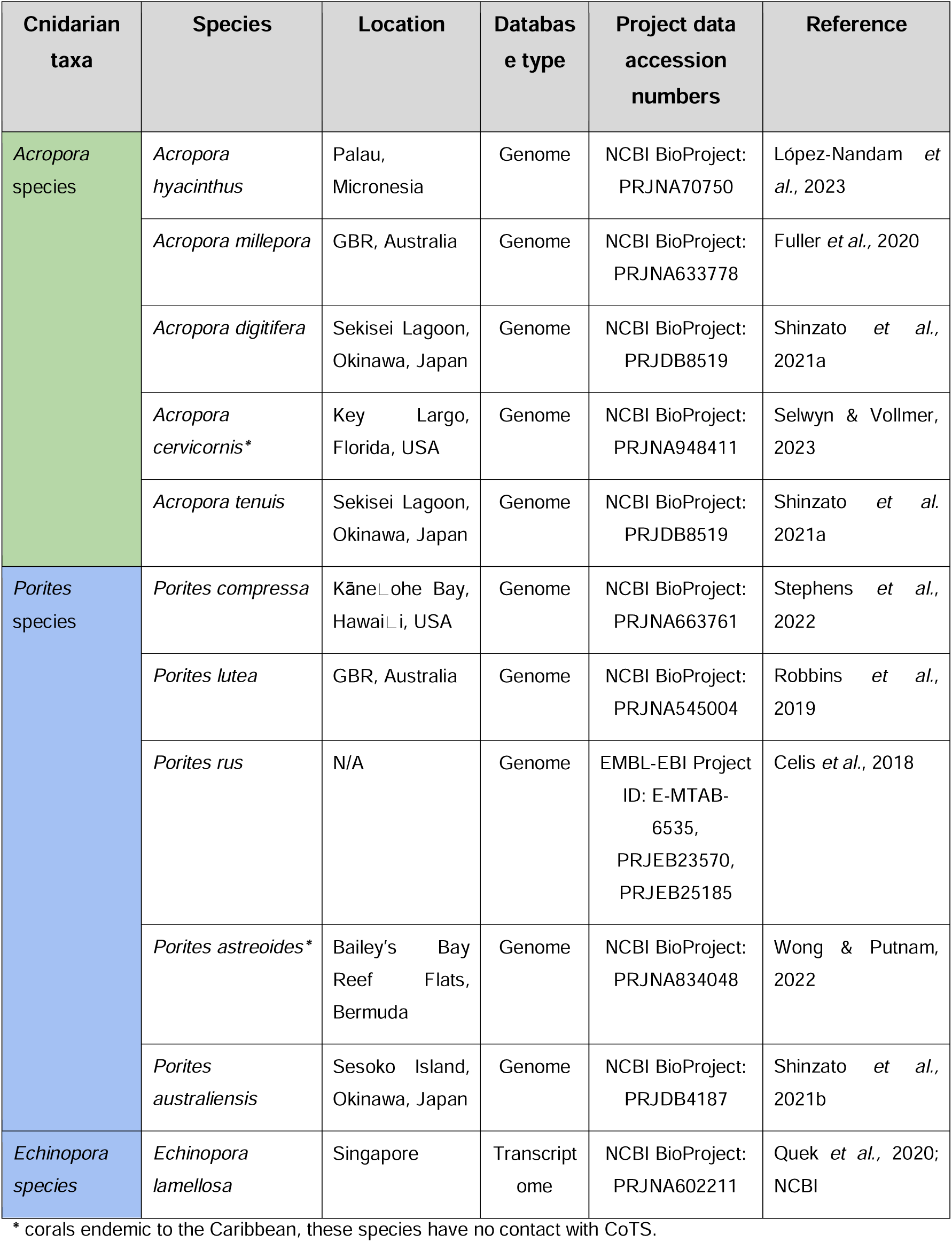
Genomic and transcriptome databases used for toxin investigations. Taxa shown in descending order from the preferred (green) prey of CoTS to the not preferred (blue).

### 2.1. Custom BLAST bioinformatics

Toxins were identified following described methods (Brinkman *et al*., 2015; Gacesa *et al*., 2015; Rachamim *et al*., 2015). Briefly, known toxin and venom peptide/protein families from cnidarians were acquired from the NCBI and UniProt databases (when available) and from the literature. The characterised toxins were then searched against InterProScan for their conserved domains and the peptides were split into groups based on their function and conserved domain sequence (**Supplemental file 1**). Toxin orthologs in each group were then aligned using MUSCLE and a conserved consensus sequence was generated (parameters: sequence similarity ≥25%; ignore gaps) in Geneious Prime (v.2024.0; https://www.geneious.com). The resulting conserved consensus sequence was used to search against the acquired cnidarian databases using BLASTp or tBLASTn searches. Toxin families with no applicable InterPro or PFAM accession numbers for conserved domains had customised conserved consensus sequences created by collecting and aligning cnidarian protein orthologs for each toxin family and BLASTing the full consensus alignment.

All reciprocal BLAST best hits (RBBHs) or relaxed RBBHs (even those with a cut-off e-value of more than 1.0e−5) from the *Acropora* and *Porites* genomes were kept. A protein redundancy score of ≥90% was used to remove any duplicate hits. The remaining hits were then BLASTed against the NCBI non-redundant and the UniProtKB/Swiss-Prot Tox-Prot databases (Jungo *et al*., 2012). Sequences with a cut-off e-value of less than 1.0e−5 to a toxin or venom constituent, a query coverage above 70 % and a relevant Gene Ontology (GO) term associated with venom synthesis and/or toxicity in the UniProtKB/Swiss-Prot Tox-Prot database were retained. The resulting sequences were then manually edited and validated. Any sequences which hit higher to a non-toxin protein family in the NCBI nr databases, or sequences with Gene Ontology terms unlikely to be related to toxins and/or venom synthesis, were removed. Top hits from both the NCBI nr database and UniProtKB/Swiss-Prot Tox-Prot database were recorded for each protein. The NCBI nr database top hit was chosen by the lowest e-value of a protein with a recognised name in the top 100 hits from a non-scleractinian coral (to avoid scleractinian paralogs). If none of the top 100 hits were proteins with a recognized name, a protein from a non-scleractinian organism with the lowest e-value in the top 100 hits was chosen. Predicted, hypothetical, uncharacterised or putative protein hits were ignored. If none of the top 100 hits were proteins with a recognized name or proteins from a non-scleractinian organism, a hit from a scleractinian coral was chosen. The top hit from the UniProtKB/Swiss-Prot Tox-Prot database was chosen based on the hit scoring the lowest e-value.

### 2.2. Discovery bioinformatics

In addition to searching the known cnidarian venom constituents and toxins, we BLASTed the whole UniProtKB/Swiss-Prot Tox-Prot database (7890 protein sequences) against the genome of each of the focal coral species to search for any toxins/venom constituents that may have been missed, and to search for putative uncharacterised toxins (Frazão *et al*., 2012; Weston *et al*., 2013; Schmidt *et al*., 2020). The BLAST hit returned for each sequence of the UniProtKB/Swiss-Prot Tox-Prot database was the hit corresponding to the lowest e-value. Redundant hits from the same genome were removed. Hits were manually edited to remove any low quality regions (e.g., duplicated regions). Hits were then re-BLASTed against the UniProtKB/Swiss-Prot Tox-Prot database and only proteins/amino acid sequences with an e-value of 1 x 10 −5 and >70% query cover were retained. Hits were also BLASTed against the InterProScan database to find conserved domains (NB: some of the translated sequences contained “*” indicating any amino acid in this position but these characters were removed from the sequence as InterProScan would not read the sequence). Gene ontology terms of the most similar toxin from the UniProtKB/Swiss-Prot Tox-Prot were also collected (**Table S11**).

### 2.3. Phylogenetics

Phylogenetic trees were constructed for the collected putative toxins from *Acropora* species, *Porites* species and *E. lamellosa* in the actinoporin, MAC-PF, CFX and SCRiP families. Our putative toxins were aligned with previously confirmed toxins in these families (**Supplemental file 1**) using MUSCLE in Geneious Prime (version 2024.0.7). Alignments were then manually edited and trimmed to the corresponding domain or conserved region (**Supplemental file 1**). Alignments were then submitted to IQ-tree (Trifinopoulos *et al*., 2016) to obtain the correct model of amino acid evolution using the corrected Akaike information criterion (AICc). The best-fit model of evolution was obtained, maximum likelihood trees were generated in IQ-tree using 1000 ultrafast bootstrap replicates and represented as SH-aLRT support (%). Trees were rooted with sister proteins from the same superfamily or homologs from distantly related species (see **Supplemental file 1** for further information). Trees were edited and visualised in Interactive Tree of Life (iTOL; version 6; Letunic & Bork, 2024).

## 3. Results

### 3.1. Custom BLAST searches *versus* discovery bioinformatics

Both our custom BLAST and discovery bioinformatics methods revealed unique proteins that were not found using alternative methods (**Table S1-11; Table 3**). This shows the importance of combining both methods for cnidarian venom discovery as only 325 of the proteins from the UniProtKB/Swiss-Prot Tox-Prot belonged to cnidarians compared with 2,409 and 2,712 from the Serpentes and Arachnida, respectively. This approach facilitated identification of toxins in the genomes of the *Acropora* and *Porites* species that were similar to those of other venomous taxa (e.g., snakes and scorpions), which our custom BLAST searches may have missed (**Table S11**). In contrast, the consensus sequences generated for our custom BLAST searches (**Supplemental file 1**) were best informed by well-studied venom proteins in Cnidaria. Thus, this method was most effective for detecting more conserved cnidarian toxins (**Table S1-10**).

**Table 3.** Total number of putative venom protein hits from discovery bioinformatics. The total number of these proteins that were found in both discovery bioinformatic methods and custom BLAST searches were then calculated. Coral taxa shown as preferred (green) and not preferred (blue) CoTS prey.

| Database | Number of venom proteins *via* discovery bioinformatics | Total number of proteins found by discovery bioinformatics which were also found by custom BLAST searches |
| --- | --- | --- |
| *Acropora cervicornis* | 86 | 13 out of 86 (15.12%) |
| *Acropora digitifera* | 53 | 12 out of 53 (22.64%) |
| *Acropora hyacinthus* | 57 | 13 out of 57 (22.81%) |
| *Acropora millepora* | 62 | 13 out of 62 (20.97%) |
| *Acropora tenuis* | 58 | 11 out of 58 (18.97%) |
| *Porites astreoides* | 37 | 12 out of 37 (32.43%) |
| *Porites australiensis* | 74 | 19 out of 74 (25.68%) |
| *Porites compressa* | 122 | 16 out of 122 (13.12%) |
| *Porites lutea* | 112 | 14 out of 112 (12.50%) |
| *Porites rus* | 59 | 15 out of 59 (25.42%) |
| *Echinopora lamellosa* | 18 | 13 out of 18 (72.22%) |

### 3.2. Venom proteins found across coral genera

The most abundant venom proteins across *Acropora, Porites* and *E. lamellosa* appeared to be neurotoxic protease inhibitors (i.e., turripeptides), neurotoxins (i.e., Kunitz-type neurotoxins), toxic enzymes (i.e., phospholipases, metalloproteinases) followed by lectins (i.e., snake lectin proteins) and pore-forming toxins (i.e., actinoporins) (**Fig. 1**; **Table S12**).

**Figure 1.**
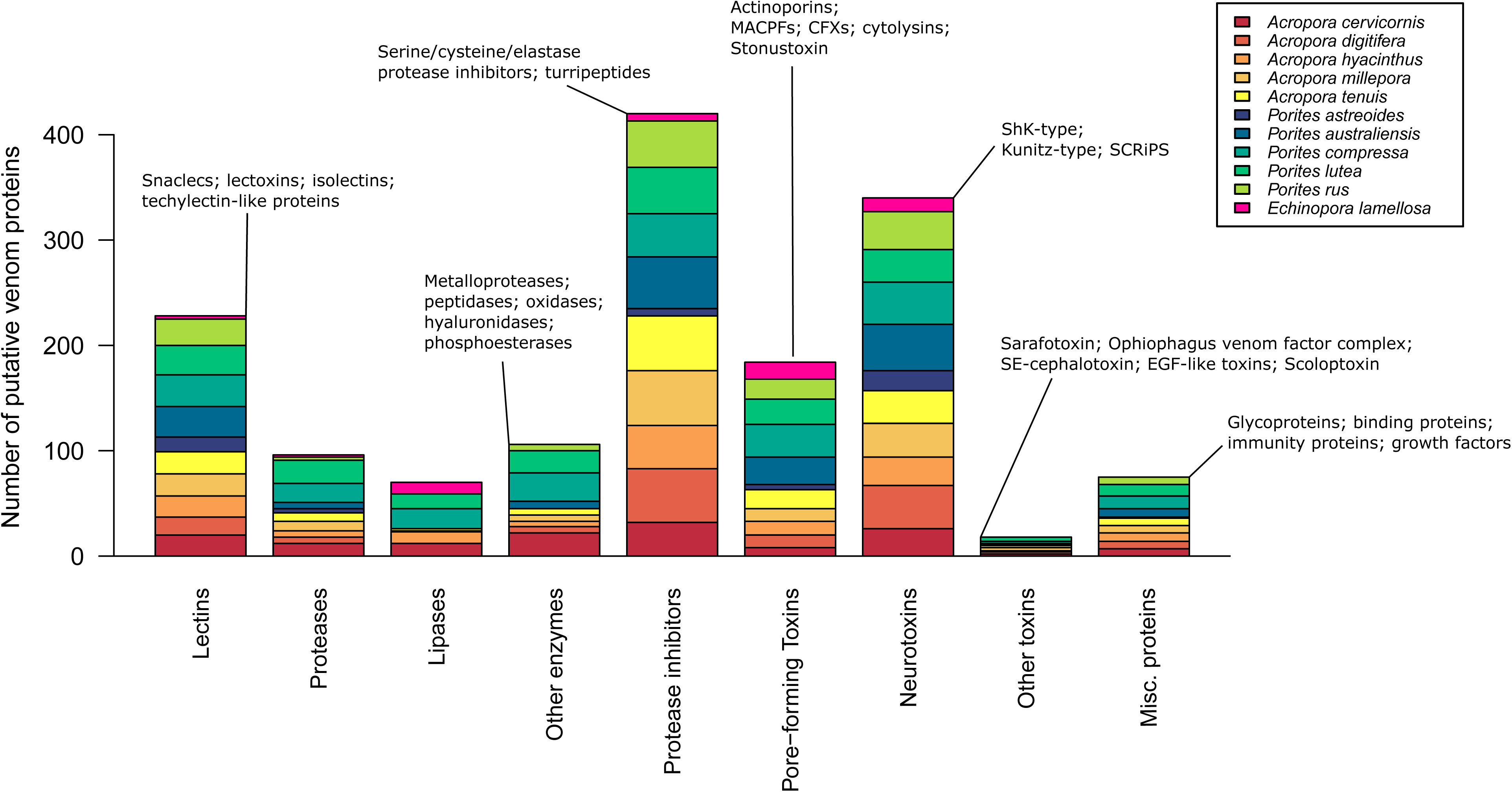
Putative coral venom protein families found across each coral species identified by custom BLAST searches and discovery bioinformatics. “Misc. proteins” (miscellaneous proteins) includes proteins that are not directly associated with venom e.g., can be found in non-venomous tissue.

*Acropora* and *Porites* species showed similar abundances of toxic lectins, proteases, lipases, other enzymes, protease inhibitors, PFTs, neurotoxins, other toxins and miscellaneous proteins across their venoms (*P* > 0.05; T-test, Mann-Whitney U test; **Fig. 1**). Despite the venoms being made up of similar abundances of protein families, we did however find several differences in the presence/absence and phylogeny of venom proteins within these protein families between *Acropora* and *Porites* species (**Fig. 2**), which we discuss in the following sections.

**Figure 2.**
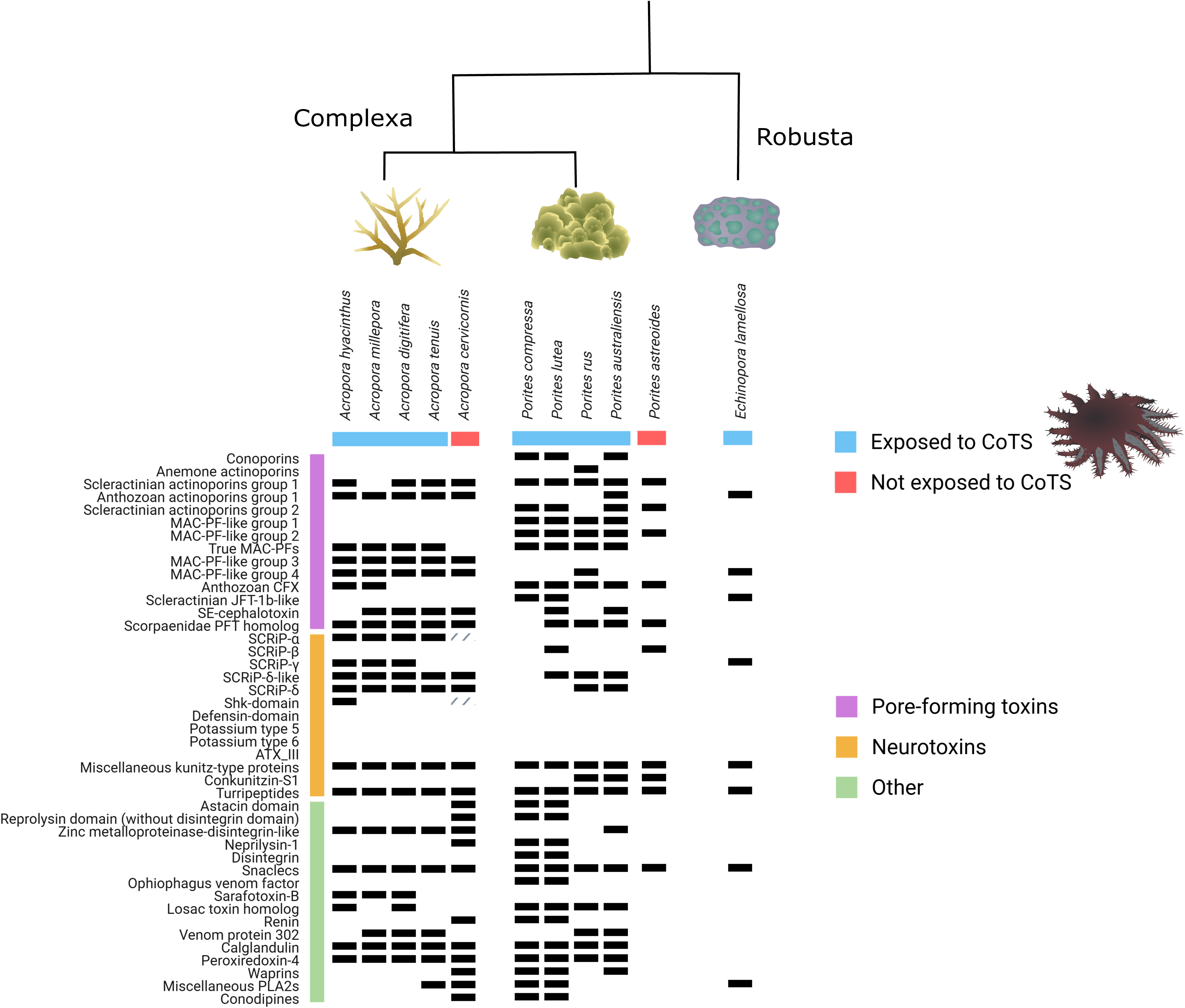
Main differences found in venom proteins across complex and robust coral genera and a coral’s exposure to CoTS (corals native to the Pacific = CoTS exposed *versus* corals native to the Caribbean = not exposed to CoTS). Black boxes represent the presence of protein found in that coral species. Actinoporin, MAC-PF, CFX and SCRiP group terminology on the y-axis are informed by phylogenetic analysis conducted in our current study (**Figs. 3-6**). Hashed boxes represent discrepancies in the data e.g., the SCRiP-α homolog was found in *A. cervicornis* in a previous study (Barroso *et al*., 2024) but no homolog was found in our current study, and we only found a protein with a partial Shk-domain in *A. cervicornis*. Graph created in BioRender (Gorman, L. (2025) https://BioRender.com/y35z919) and figure edited in InkScape. Images of CoTS and coral species downloaded from the media library at the Integration Network University of Maryland Centre for Environmental science (https://ian.umces.edu/media-library/).

#### 3.2.1 Cytolysins and pore-forming toxins (PFTs)

In total, 49 actinoporins/cytolysins were found across the *Acropora* and *Porites* species investigated, while *E. lamellosa* contained four putative actinoporins (**Table S1, S11, Fig. 3**). Phylogenetic analysis of the 53 actinoporins we discovered (**Table S1**) showed that our putative actinoporins from *Acropora* species, *Porites* species and *E. lamellosa* were present across five of the seven actinoporin phylogenetic groups (**Fig. 3**). None of our actinoporins grouped with the MAC-PF toxins or hydralysins we included in our actinoporin tree, which both made distinct sister clades with 100% bootstrap support (**Fig. 3**). Three actinoporin hits exclusively from *Porites* species (one hit from each of *P. compressa*, *P. lutea* and *P. australiensis*) clustered with tereporins/conoporins with 100% bootstrap support in the tree and also hit to tereporins/conoporins when BLASTed against the UniProtKB/Swiss-Prot Tox-Prot database (**Table S1, Figs. 2, 3**). It should be noted however, that only the hit from *P. compressa* matched the query cover quality criteria set (>70%). That said, we included all three putative tereporin/conoporin homologs from the three *Porites* species, as they all had low e-values (<1 x 10 ^-5^; **Table S1**). One putative PFT sequence form *Porites rus* also showed a distinct phylogenetic relationship, being the only PFT we discovered across the 11 coral species that clustered with haemolytic anthozoan PFTs (‘Anemone actinoporins’; **Fig. 3**). Despite this, none of our coral species’ putative actinoporins were related to the confirmed scleractinian haemolytic actinoporin, Δ-Pocilopotoxin-Spi1 from *Stylophora pistillata* (Ben-Ari *et al*., 2018; **Fig. 3**).

**Figure 3.**
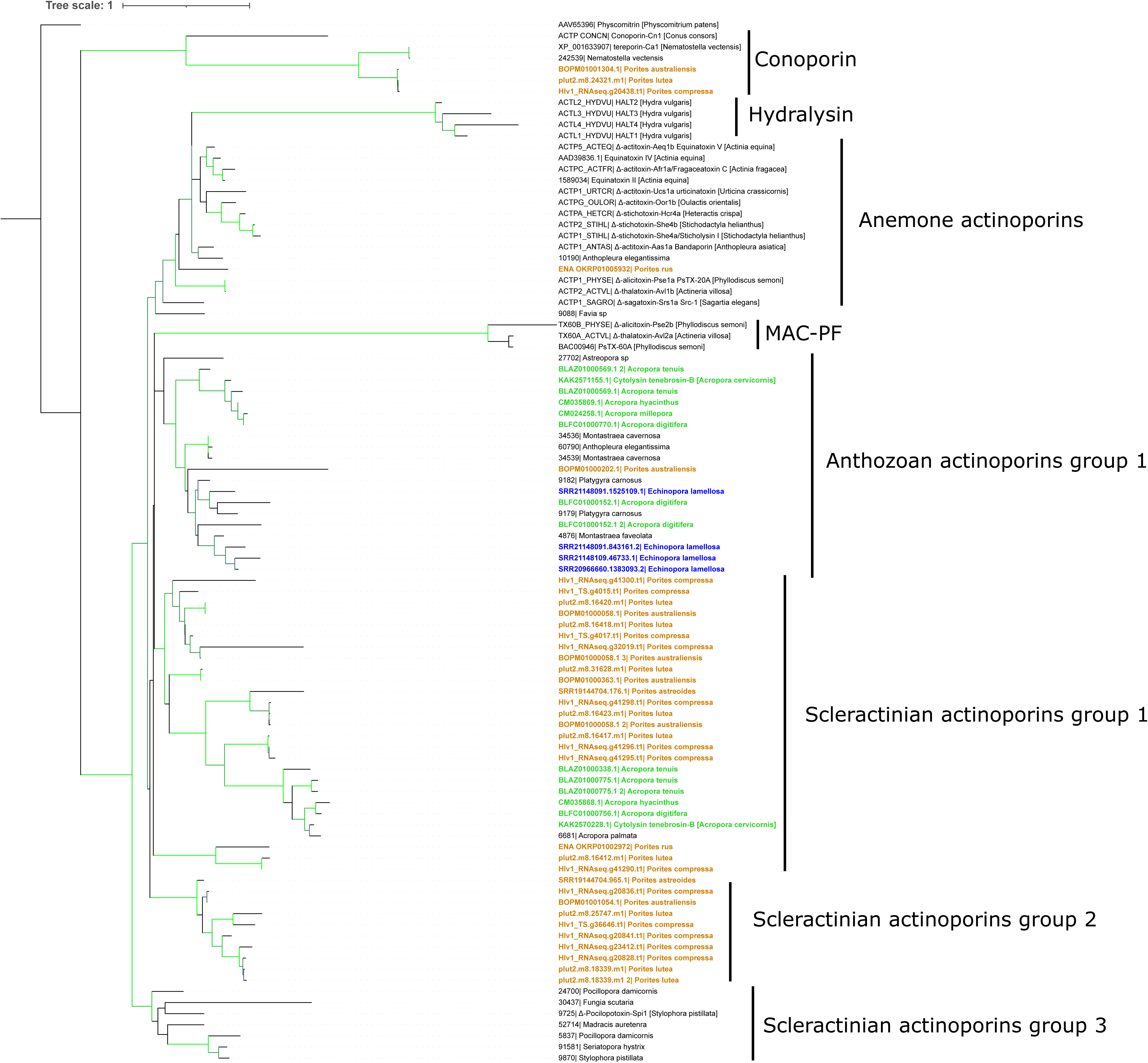
Maximum likelihood tree of actinoporins across Cnidaria. Putative actinoporins from the current study are denoted in green (*Acropora* species), brown (*Porites* species) and blue (*Echinopora lamellosa*). Model of evolution and tree constructed in IQ tree (Trifinopoulos *et al*., 2016) and tree visualised and edited in Interactive tree of life (iTOL; Letunic & Bork, 2024). Only branches with bootstrap support >70% are coloured in green.

Only four hits were returned for the hydralysin consensus sequence (**Table S2**) and all hits had high e-values (>1 x 10 ^-5^) and in addition contained the actinoporin anemone cytotox domain, suggesting that these hits were in fact just actinoporins that were distantly related to hydralysins. In addition, another superfamily of PFTs, the Membrane Attack Complex/Perforin (MAC-PF) toxins were found across *Acropora* and *Porites* species and *E. lamellosa,* with 73 total MAC-PFs being found across all species investigated and being spread across five distinct phylogenetic MAC-PF groups (**Table S3, S10, Fig. 4**).

**Figure 4.**
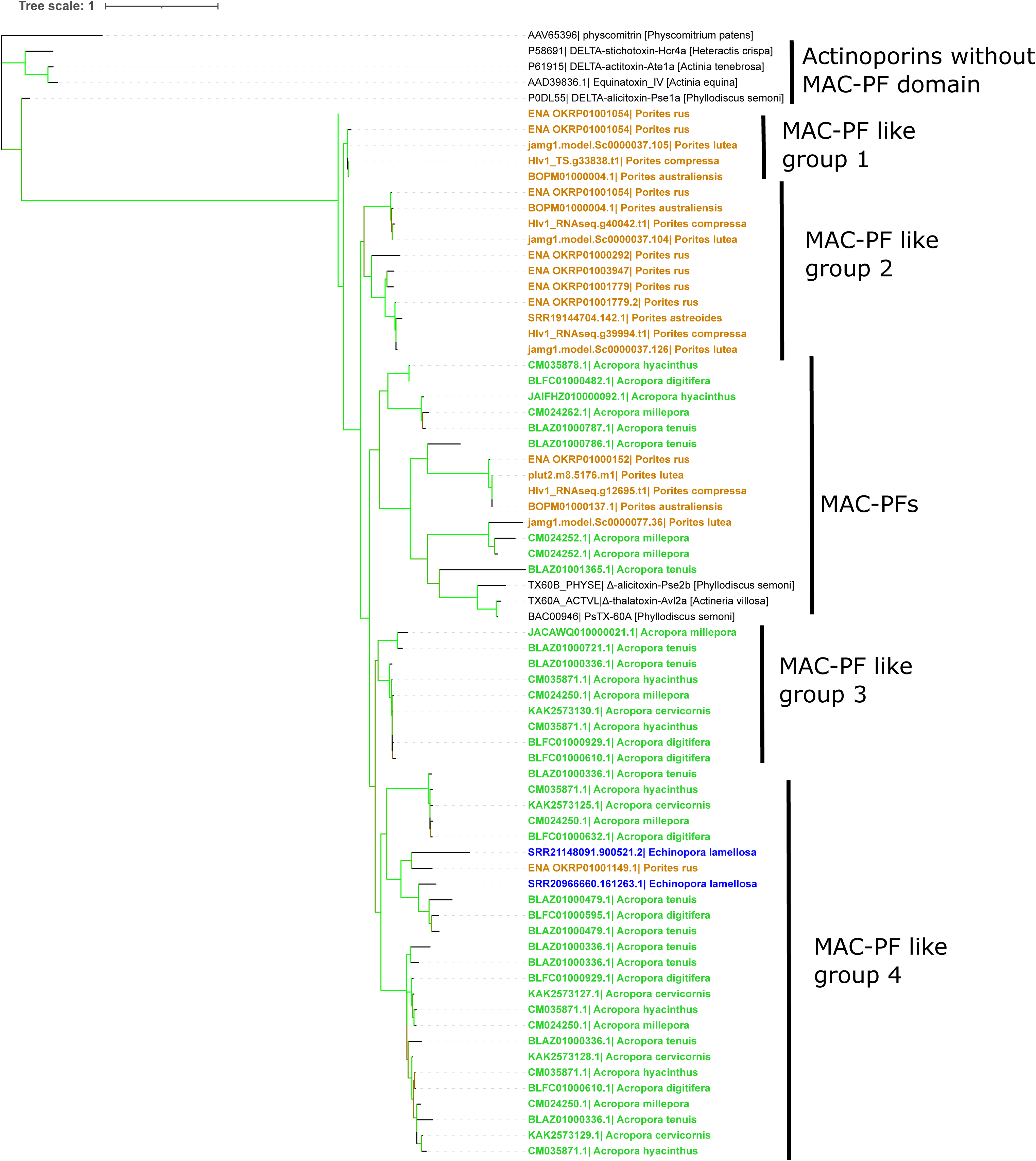
Maximum likelihood tree of Membrane Attack Complex/Perforin (MAC-PF) toxins and their homologs. Sea anemone MAC-PFs were gathered from Frazão *et al*., (2012) and aligned with putative MAC-PFs from *Acropora* species (green), *Porites* species (brown) and *Echinopora lamellosa* (blue). Model of evolution and tree constructed in IQ tree (Trifinopoulos *et al.,* 2016) and tree visualised and edited in Interactive tree of life (iTOL; Letunic & Bork, 2024). Branches with >70% bootstrap support shown in light green.

#### 3.2.2. *Chironex fleckeri*-like toxins (CFXs)

Custom BLAST searches used a consensus alignment from eight CFX proteins. Only two *Acropora* species (*A. millepora* and *A. hyacinthus*) returned hits to CFXs, each containing only one putative CFX. Contrastingly, all *Porites* species included in this study returned hits to CFXs, ranging from one putative CFX in *Porites astreoides* to 15 in *Porites compressa* (**Table S4**). Altogether 46 putative CFX proteins were returned for the *Acropora* and *Porites* species investigated and none of these proteins contained any recognised conserved domains (**Table S4**). No CFX hits were returned from the *E. lamellosa* transcriptome. To investigate whether this result was due to too many ambiguous amino acid residues in the consensus CFX alignment (amino acid residues corresponding to “X”; **Supplemental file 1**), we also BLASTed each individual CFX used in the consensus alignment against the *E. lamellosa* transcriptome. This method returned one hit to CfTX-B from *C. fleckeri* (UniProt: T1PQV6; **Table S4**).

The putative CFX-like toxins from *Acropora* species, *Porites* species and *E. lamellosa* clustered across two phylogenetic groups (**Fig. 5**). One group harboured several putative CFX proteins from *Porites* species and one from *E. lamellosa* and these putative CFXs grouped with jellyfish JFT-1b toxins such as *C. fleckeri* toxins B and A (86.6% bootstrap support; **Fig. 5**). This same group clustered with the Cry-like toxins from scleractinian corals *Stylophora pistillata* and *Madracis auretenra* (81.7% bootstrap support). The second group contained all *Acropora* CFXs and the majority of putative *Porites* CFX-like proteins. This second group was an anthozoan-specific JFT clade, being a phylogenetically distinct sister clade to true JFTs (86.4% bootstrap support; **Fig. 5**). This group contained CFX-like toxins from other anthozoans e.g., *Gorgonia ventalina, Exaiptasia pallida* and *Eunicella cavolini* (**Fig. 5**). None of the putative CFXs from *Acropora* species, *Porites* species or *E. lamellosa* grouped with JFT-1a,c or JFT-2a,b toxins (Klompen *et al.,* 2021; **Fig. 5**).

**Figure 5.**
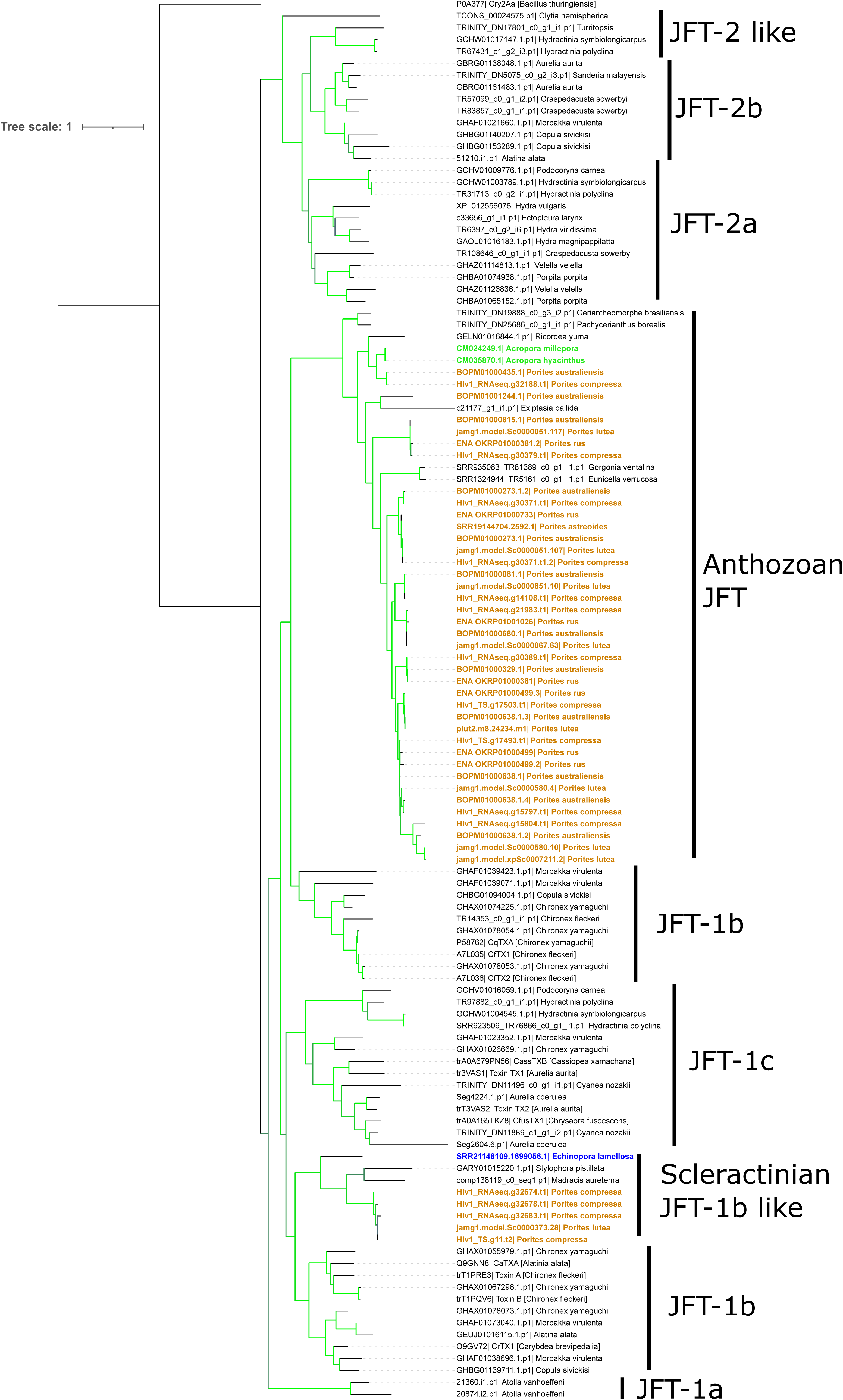
Maximum likelihood tree of jellyfish toxins (JFTs) and their homologs. JFTs gathered from Klompen *et al*. (2021) and aligned with putative CFXs from *Acropora* species (green), *Porites* species (brown) and *Echinopora lamellosa* (blue). Model of evolution and tree constructed in IQ tree (Trifinopoulos *et al*., 2016) and tree visualised and edited in Interactive tree of life (iTOL; Letunic & Bork, 2024). Groupings named as in Klompen *et al*. (2021). Branches with >70% bootstrap support shown in light green.

#### 3.2.3. Neurotoxins

For our custom BLAST bioinformatics, we identified neurotoxins from a range of cnidarians and divided them into groups based on their conserved domain regions identified by InterProScan (ATX_III; defensin-like; ShK; SCRIPs; Kunitz-type; kazal turripeptides; and potassium type 5 and 6 neurotoxins; **Supplemental file 1**). We found putative neurotoxins with similarities to kunitz-domain neurotoxins (**Table S5, S10**); SCRiPS (**Table S6, S11**); ShK-domain containing neurotoxins (**Table S7, S11**); turripeptides containing the kazal 1 and 2 domains (**Table S8, S11**) in our custom and discovery bioinformatics methods. This contrasts to a previous study that found no evidence of Shk-like neurotoxins in scleractinian corals (Schmidt *et al*., 2019). However, most of the possible Shk-domain neurotoxins we found were classed as low quality proteins, with only one putative Shk protein from *Acropora hyacinthus* meeting the quality criteria in custom BLAST searches (**Table S7**) and only one protein (KAK2558191.1) from *Acropora cervicornis* containing a partial Shk domain in the discovery bioinformatics (**Table S11**). No putative neurotoxins containing the ATX_III or defensin-like domain were found in *Acropora* species, *Porites* species or *E. lamellosa*, nor were any hits returned for potassium type 5 or type 6 neurotoxins (**Fig. 2**).

SCRiPS were found across *Acropora* and *Porites* species and *E. lamellosa*, returned from both custom BLAST searches and discovery bioinformatics (**Tables S4, S6, S11**). Our SCRiP phylogenetic data shows that the majority of *Porites* and *Acropora* species harboured putative SCRiPS that grouped with (“*SCRiP-*δ*-like”*) or alongside the *SCRiP-*δ clade (“*SCRiP-*δ*”*; **Figs. 2, 6**). The SCRiP-α clade was only present across *Acropora* species (*A. tenuis*, *A. digitifera*, *A. hyacinthus* and *A. millepora*; **Fig. 6**), agreeing with a previous study that concluded the SCRiP-α clade was only present in Acroporidae corals (Barroso *et al*., 2024).

**Figure 6.**
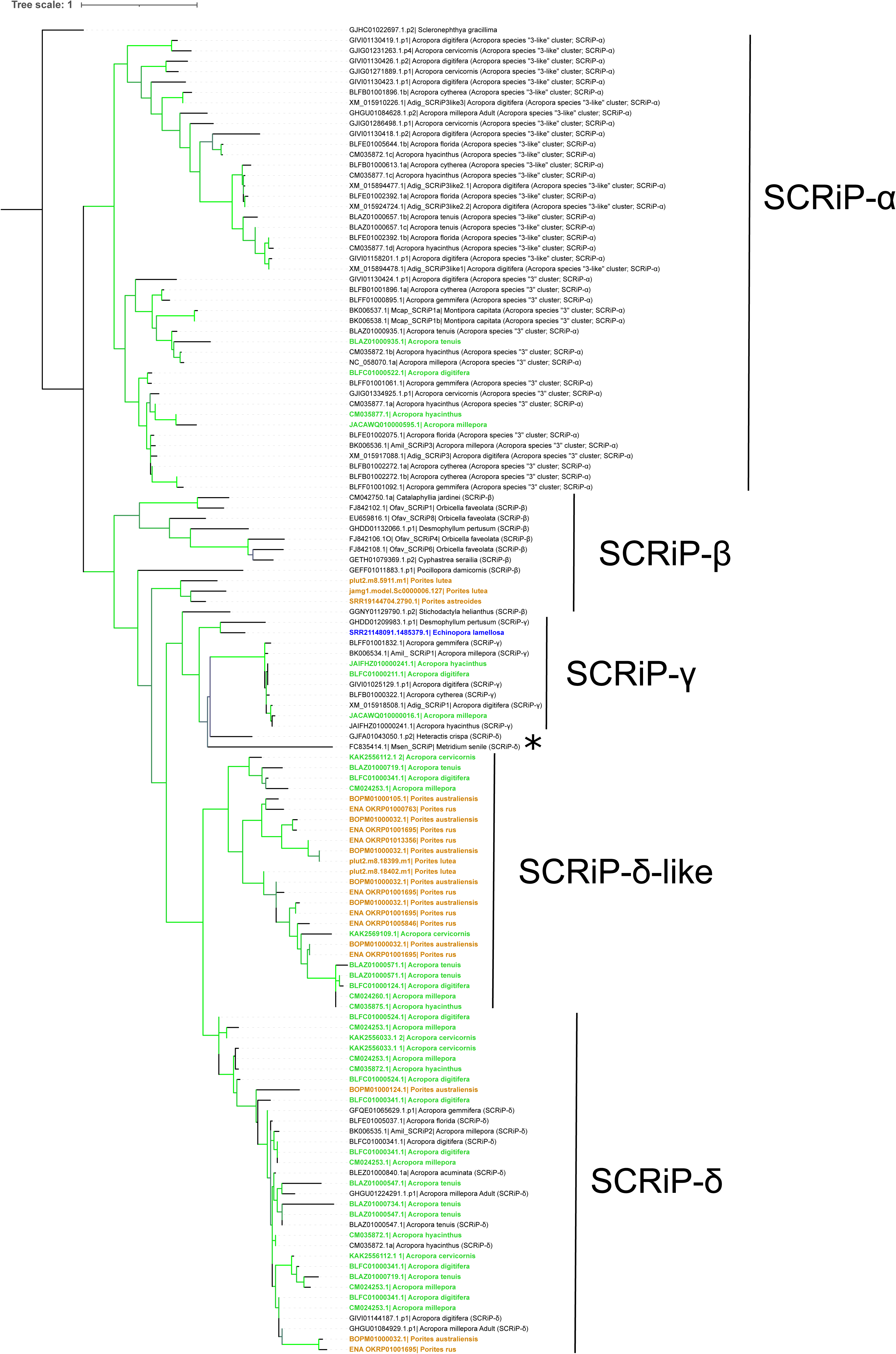
Small Cysteine-Rich Proteins (SCRiPs) maximum likelihood tree. Putative SCRiP homologs from this current study are denoted in green (*Acropora* species), brown (*Porites* species) and blue (*Echinopora lamellosa*). Other sequences were taken from Barroso *et al*. (2024). Groups named accordingly with those named in Barroso *et al*. (2024). Model of evolution and tree constructed in IQ tree (Trifinopoulos *et al*., 2016) and tree visualised and edited in Interactive tree of life (iTOL; Letunic & Bork, 2024). Branches with >70% bootstrap support shown in light green. Astericks (*) represents tree topology that differs from Barroso *et al*. (2024).

In addition to the absence of SCRiP-α in *Porites*, *Porites* species also did not harbour a SCRiP-[homolog, contrasting to both *Acropora* species (*A. digitifera*, *A. hyacinthus* and *A. millepora*) and *E. lamellosa* (**Fig. 6**). *Porites* species (*P. lutea* and *P. astreoides*) did however harbour homologs to the SCRiP-β, whilst *Acropora* species and *E. lamellosa* did not.

In addition, homologs to the neurotoxic enzymes - acetylcholinesterase (AChE) and phospholipase A2s were found by discovery bioinformatics across *Acropora* and *Porites* species. Phospholipases are regarded as venom allergen proteins, of which we also found homologs of other venom allergen proteins (all containing the cysteine-rich secretory protein, antigen 5, and pathogenesis-related 1 protein (CAP) domain; **Table S12**) as well as homologs to hyaluronidase (all containing the Glyco_hydro_56 domain; **Table S12**) across the genomes of the *Acropora and Porites* species. One *Acropora* and two *Porites* species (*A. cervicornis*, *P. lutea* and *P. compressa*) harboured PLA2 venom allergens homologs to those of cone snails (conodipines) (**Fig. 2**).

#### 3.2.5. Metalloproteases

The zinc-binding motif (HExxHxxxxxH) sequence of metalloproteases is highly conserved in cnidarians (Hwang *et al*., 2022) and was present in the consensus sequence in our custom BLAST searches (**Supplemental file 1**). Metalloproteinase results from custom BLAST searches were different to the results obtained from the discovery bioinformatics. For example, hits returned by discovery bioinformatics contained eight metalloproteinase domains not found by our custom BLAST searches (**Table S11, S12**). Contrastingly, all returned hits from the custom BLAST searches contained either the peptidase M10 or/and the hemopexin domains, albeit with low query covers and e-values (**Table S10**). Therefore, due to them being classed as low quality results, hits to the peptidase M10 and hemopexin were not included in the results (**Table S12**).

#### 3.2.6. Other associated proteins

Discovery bioinformatics revealed that venom proteins containing the lectin C domain (PFAM ID:PF00059) were widespread throughout all the venoms that we investigated. These proteins mainly corresponded to snake venom snaclec (**<u>SNA</u>**ke **<u>C</u>**-type **<u>LEC</u>**tins) proteins (**Figs. 1**, **2, Tables S11, S12**). In addition to snaclecs, other toxic lectin homologs were present across *Acropora* and *Porites* species such as lectoxin (derivatives present across all *Acropora* and *Porites* species), isolectin (Sp-CL4), and the galactose-specific lectin Nattectin (**Table S11**).

Homologs to unique toxins/proteins with unknown functions during envenomation were also present across *Acropora* and *Porties* species e.g., renin; a homolog to a venom peptide from scorpion venom (venom peptide 302; UniProt: P0CJ14), containing an insulin-like growth factor-binding protein (IGFBP) domain; homologs to trehalase; and, certain cysteine-rich venom proteins e.g., Mr30 (Qian *et al*., 2008) (**Tables S11, S12**). Contrastingly, several unique toxin homologs with more well-known functions, e.g., SE-cephalotoxin, caterpillar toxin losac (*Lonomia obliqua* Stuart factor activator) and Stonustoxin homolog, were found in *Acropora* and *Porites* species putative venoms (**Table S11, Fig. 2**).

### 3.3. Differences in venom proteins between *Acropora* and *Porites* species

Despite most venom protein families being found across all coral species we investigated, some proteins displayed species-specific affiliations (**Fig. 2**). For instance, only certain *Porites* species contained the gastropod tereporin/conoporin toxin (*P. australiensis*, *P. lutea* and *P. compressa*) and homologs to the snake venom protein - ophiophagus venom factor (*P. compressa* and *P. lutea*). The proteins homologous to the ophiophagus venom factor in *Porites* contained a unique protein domain structure (8-9 distinct protein domains) not observed in the rest of the venom protein hits (**Table S12**). A homolog to the cardiotoxin sarafotoxin-B from the snake *Atractaspis engaddensis* was found across three *Acropora* species (*A. digitifera*, *A. millepora* and *A. hyacinthus*) but absent in *Porties* species (**Table S11, Fig. 2**).

## 4. Discussion

Our study has shown the efficiency of using both custom BLAST searches in addition to a discovery bioinformatics method to discover both conserved and novel putative venom proteins across cnidarian venoms. Although our study is a non-targeted bioinformatics study utilising publicly available genomes and transcriptomes, our results are similar to those of venom-targeted laboratory studies of cnidarian species. For instance, we found that the most abundant venom proteins present across both *Acropora* and *Porites* species were protease inhibitors (e.g., turripeptides), neurotoxins (e.g., Kunitz-type neurotoxins), toxic enzymes (e.g., phospholipases, metalloproteinases) followed by lectins (e.g., snake lectin proteins) and pore-forming toxins (e.g., actinoporins). The identification of these venom protein families agrees with previous studies, including venom-targeted laboratory studies, on hydrozoans (Jaimes-Becerra *et al*., 2019; Hernández-Elizárraga *et al*., 2023), sea anemones (Castañeda & Harvey, 2009; Frazão *et al*., 2012; Orts *et al*., 2013; Jaimes-Becerra *et al*., 2019; Finol-Urdaneta *et al*., 2020; An *et al*., 2022), jellyfish (Weston *et al*., 2013), scleractinian corals (Schmidt *et al*., 2019), staurozoans (Jaimes-Becerra *et al*., 2019), cone snails (Safavi-Hemami *et al*., 2014) and snake venoms (Tasoulis & Isbister, 2023), that have all shown these protein families make up the majority of venom proteins from these species. Although we found large overlaps between the protein families present across our coral species, we did find several notable differences. For instance, *Porites* species contain homologs to both a cone snail neurotoxin (conkunitzin-S1) and cone snail PFTs (conoporins) which were absent in all *Acropora* species and *E. lamellosa* we investigated*. Porites* species and *E. lamellosa* also harboured putative CFX homologs that clustered with jellyfish whereas putative *Acropora* CFXs clustered with anthozoans. Additionally, we found the presence of certain snake venom proteins that were present in *Acropora* species (sarafotoxin) that were absent in *Porites* species and *vice versa* (ophiophagus venom factor).

### *Porites* species show venom homologs similar to those of cone snails, jellyfish and snakes which are absent in *Acropora* species

Unsurprisingly, we found pore-forming toxin (PFT) homologs (actinoporins, MAC-PFs, CFXs) in all the species we studied, as PFTs represent a major class of toxins in cnidarian venoms (Frazão *et al*., 2012; Rachamim *et al*., 2015). Although all the coral species we investigated contained these classes of proteins, we observed interesting differences. For example, only *Porites* species (*P. australiensis*, *P. lutea* and *P. compressa*) contained homologs to tereporin/conoporin toxins, and only *Porites* species (*P. compressa* and *P. lutea*) and *E. lamellosa* harboured CFX homologs that grouped with the jellyfish JFT-1b CFXs. Other studies have also found conoporin- and CFX-like proteins in cnidarian venoms (Jaimes-Becerra *et al*., 2019; Klompen *et al*., 2020; Klompen *et al*., 2021), although it is currently debated whether anthozoans truly contain CFX-like toxins or rather a actinoporin sister family related to CFXs, with no CFX homologs isolated from anthozoan venoms (Klompen *et al*., 2021). Nevertheless, the absence of conoporin homologs in all *Acropora* species we investigated, in addition to the absence of JFT-1b CFX homologs in *Acropora* species is interesting, especially when considering that these homologs were present in *Porites* species, the non-preferred prey of CoTS. Cone snails and jellyfish are known to contain sublethal and lethal toxin repositories (Nisa *et al*., 2021; Ratibou *et al*., 2024) and as *Porites* species contain homologs of these proteins (conoporins and JFT-1b CFXs), it may imply that *Porites* species harbour toxins that are more effective at causing injuries to CoTS and therefore deterring CoTS predation. For instance, conoporins and tereporins are potent PFTs that target the nervous system inducing paralysis in victims (Turner *et al*., 2018). The relationship between these phylogenetic differences and specific PFT homologs and their bioactive role in predator defence remains unknown. However, a study comparing haemolytic activity of corals (Ben-Ari *et al*., 2018), showed species-specific differences. For instance, *Stylophora pistillata*, whose venom contains the haemolytic actinoporin Δ-Pocilopotoxin-Spi1, elicited a haemolytic activity on human erythrocytes that was 600 times greater than that of other scleractinian venoms (Ben-Ari *et al*., 2018). This emphasises that a coral’s repertoire of PFTs likely contributes to the bioactivity of its venom and therefore its efficacy on predators/prey.

Even though all corals we investigated harboured homologs to kunitz-type neurotoxins, only *Porites* species (*P. astreoides*, *P. australiensis* and *P. rus*) contained a homolog to the cone snail kunitz-protein, conkunitzin-S1 (Bayrhuber *et al*., 2005). A previous study also found a homolog to conkunitzin-S1 in the transcriptome of the CoTS predator *Charonia tritonis* (Zhang *et al*., 2022). Furthermore, the authors synthesised the *C. tritonis* conkunitzin-S1 homolog (Ct-kunitzin) and investigated its bioactivity against mice and CoTS in laboratory assays (Zhang *et al*., 2022). In mice, Ct-kunitzin suppressed pain recognition and reduced their locomotory activity, whilst in CoTS, Ct-kunitzin destroyed the tube feet (Zhang *et al*., 2022). In the black mamba snake venom (*Dendroaspis polylepis*) kunitz-type toxins (known as dendrotoxins) are amongst the most potent venom components, exhibiting exceptionally high lethality (Laustsen *et al*., 2015). The potential for a conkunitzin-S1 homolog in *Porites* to elicit similar bioactivity as seen from Ct-kunitzin, and contribute to a venom defence more efficient against CoTS predation warrants further investigation.

We also found a homolog to the snake venom protein, ophiophagus venom factor (OVF) from the king cobra *Ophiophagus hannah* only present in *Porites* species (*P. compressa* and *P. lutea*). OVF (also known as the cobra venom factor (CVF)) is a non-lethal depletion factor of the complement system (intrinsic part of the immune system), with a high homology to mammalian complement C3 and shares functional similarity (Vogel & Fritzinger, 2010). At the site of envenomation, OVF releases anaphylatoxins (e.g., C5a) in humans, increasing vasodilation, reducing blood pressure and eliciting vascular permeability that in turn facilitates the spread of venom and leads to the paralysis and death of the prey (Vogel & Fritzinger, 2010; Zeng *et al*., 2012; Tambourgi & van den Berg, 2014). Homologs to CVF have also been found in other cnidarian venoms (Brinkman *et al*., 2015; Jaimes-Becerra *et al*., 2019). The potential for the OVF homolog in *Porites* to function similarly to its counterpart in snake venom - facilitating more rapid and/or efficient spread of the venom, merits future investigation. The absence of this protein in *Acropora* species venom is intriguing, and may result in *Acropora* species venom not spreading as successfully in CoTS upon envenomation compared to venom from *Porites* species.

Although we did find *Porites* species to contain more venom proteins that were absent from *Acropora* species, we did find one venom protein in *Acropora* species (*A. digitifera*, *A. millepora* and *A. hyacinthus*) that was absent from all *Porites* species, Sarafotoxin-B. Sarafotoxin-B is a cardiotoxic peptide from venom of the snake *Atractaspis engaddensis,* that elicits vasoconstriction through its binding to endothelin receptors on atrial and brain membranes, and activating the hydrolysis of phosphoinositides, leading to impairment of the left ventricular function of the heart (Kloog *et al*., 1988; Mahjoub *et al*., 2015). If this toxin has a similar function in *Acropora* venom, the bioactivity against CoTS would need to be investigated as endothelin receptors are only present in vertebrates (Davenport *et al*., 2016).

### Both *Acropora* and *Porites* species contained venom homologs to unique toxins

In addition to identifying cnidarian PFTs, our bioinformatics analysis discovered a myriad of other homologs present across both *Acropora* and *Porites* species to unique venom toxins and their conserved domains. Notably, a SE-cephalotoxin homolog, a Scorpaenidae PFT homolog containing the Stonustoxin_helical (PFAM ID: PF21109) and Thioredoxin_11 (PFAM ID: PF18078) domains, and a homolog caterpillar toxin called losac (Lonomia obliqua Stuart factor activator) (**Table S11**). Although these toxins have unique biochemical structures, all of them have shown to elicit potent haemolytic and/or neurotoxic activities during envenomation (Ueda *et al.,* 2006, 2008; Alvarez-Flores *et al*., 2011; Malacarne *et al*., 2018; Gonçalves & Costa, 2021; Maru *et al*., 2021). SE-cephalotoxin initially showed no homology to other confirmed toxins when it was first identified (Ueda *et al.,* 2008), however more recent studies have shown venom proteins with sequence similarity to SE-cephalotoxin present in stingray venom (Silva *et al*., 2018), emphasising that toxins with this structure may be more widespread in venomous marine taxa than originally thought. The plethora of homologs to unique toxins with diverse bioactivity and structures present across both *Acropora* and *Porites* species, highlights the extensive venom arsenal that these species can potentially deploy during CoTS predation. However, it is not known whether all or only a subset of these toxins are utilised against CoTS during predation, and whether their deployment is phenotypically plastic with regard to environmental cues.

In addition, we found that homologs to snaclec proteins comprised the majority of the toxic lectins found across all our coral species. Snaclecs are non-enzymatic snake venom proteins that bind in a Ca^2+^-dependent manner to mono- and oligosaccharides (Ogawa *et al*., 2005). In the presence of Ca^2+^, Snaclecs facilitate a diverse array of functions as they selectively target different membrane receptors, coagulation factors and proteins involved in hemostasis (Weis *et al*., 1998; Arlinghaus & Eble, 2012). Due to the diverse function of snaclecs despite their conserved homology (Weis *et al*., 1998), it would be difficult to suggest a possible function for snaclecs homologs in scleractinian venom without further functional and bioactivity studies.

### Neurotoxins and toxic enzymes were widespread throughout the coral species studied

Unsurprisingly, enzymes and neurotoxins were a major feature of *Acropora* species, *Porites* species and *E. lamellosa* venoms (e.g,. acetylcholinesterase, hyaluronidase, PLA2, metalloproteinases, turripeptides, SCRiPS and kunitz-type neurotoxins). Enzymes are an important component of venoms, functioning not only in direct toxicity but also in the spread of the venom, activating other venom components and helping to preserve the venom (Gonçalves & Costa, 2021; Delgado-Prudencio *et al*., 2022). All the metalloproteinase hits we retrieved from the custom BLAST searches were low quality when BLASted to the UniProt Tox Prot database. Whether this is truly due to the proteins being low quality or the fact that these families of metalloproteases have diverged significantly in scleractinians warrants future investigation. This latter hypothesis is a potential possibility due to our BLAST searches against the NCBI nr database returning significant hits to other species metalloproteases e.g., anthozoans and echinoderms (**Table S10**). Indeed, harbouring a large repertoire of different families of neuroactive peptides would allow efficient paralysis of prey and/or predators.

Scleractinian neurotoxins SCRiPS were found across all the species we investigated and our phylogenetic data of *Acropora* SCRiPS agrees with that of a recent study (Barroso *et al*., 2024). However, this same study did not find homologs to SCRiPS in *P. rus* and *P. australiensis* genomes or a *P. lutea* transcriptome (Barroso *et al*., 2024), contrasting to our results from both our custom BLAST and discovery pipelines. We found large expansions of SCRiPS homologs in *Porites* and *Acropora* species in the SCRiP-δ and SCRiP-δ-like clades. As SCRiP-δ was the only SCRiP clade detected in sea anemones and had the largest purifying selection (toxin potency preserved after initial expansion) and a SCRiP-δ from *A. millepora* had the highest potency versus a *SCRiP-*α (Jouiaei *et al*., 2015), it was hypothesised that this may be the main SCRiP functioning in predator defense (Barroso *et al*., 2024). Our data would agree with this, emphasised by the large expansion of the SCRiP-δ families in both *Porites* and *Acropora* species, emphasising that SCRiP-δ must play an important role in the biology of these corals. Future studies should investigate the data provided by this study and the previous studies on SCRiPS (Jouiaei *et al*., 2015; Barroso *et al*., 2024) to investigate the role and function of SCRiPS (especially proteins from the SCRiP-δ family) in predator defence of coral species against CoTS.

### We found proteins in all coral species with unknown function and those not involved in envenomation

In addition to venom protein homologs with known functions in venom, we also identified several homologs to venom proteins with functions that remain uncertain. For instance, a homolog to renin was present across both *Acropora* and *Porites* species. In snake venom, renin is an aspartic protease that cleaves angiotensinogen into angiotensin-1 (Wilkinson *et al*., 2017). Although the function of renin in snake venom is unconfirmed, it is hypothesised to contribute to hypertension in humans following envenomation (Wilkinson *et al*., 2017). Additionally we found the presence of a venom protein containing the IGFBP domain across several *Acropora* and *Porites* species (venom protein 302), agreeing with other studies on cnidarian venoms (Brinkman *et al*., 2015; Liao *et al*., 2019; Klompen *et al*., 2022). Proteins containing the IGFBP domains have also been found in scorpion (Ward *et al*., 2018) and spider venoms (Kuhn-Nentwig *et al*., 2011). However, it is unclear whether IGFBP contributes to envenomation as IGFBPs are also expressed in non-venom related tissues of venomous species (Kuhn-Nentwig *et al*., 2011; Valdez-Velázquez *et al*., 2020). In non-venomous tissue, IGFBPs function as carrier proteins that regulate the transport, turnover and distribution of insulin growth factors (IGFs) (Duan & Xu, 2005). IGFs are polypeptides that are involved in cellular proliferation, differentiation and homeostasis (Ranke & Elmlinger, 1997). Similarly to IGFBPs, we also found the presence of the enzyme trehalase, which is also found in non-venomous tissue, whose function is to break down the blood sugar trehalose, and therefore, trehalase regulates energy metabolism and glucose generation (Shukla *et al*., 2015). Thus, the function of these proteins may be to disrupt the metabolic equilibrium in victims, as shown in other animal venoms post envenomation e.g., parasitoid wasps (Mrinalini *et al*., 2015) and snakes (Matkivska *et al*., 2023).

In addition to proteins with unknown functions, we also found the presence of proteins hypothesised to function in the venom cell but not directly in the venom itself e.g., calglandulin and peroxiredoxin 4. Calglandulin exports toxins out of the cell and into the venom gland in snakes (Zhang *et al*., 2006) whilst peroxiredoxin 4 is an antioxidant enzyme hypothesised to lead to the diversification of toxins in the venom gland of snakes (Calvete *et al*., 2009) (**Table S11**). We also found homologs to waprins (omwaprin-a and -b in *P. lutea*, *P. compressa*, *P. australiensis* and Waprin-Phi3 in *A. cervicornis*). Waprins are snake venom peptides that show protease inhibitor activity as well as significant antimicrobial activity (Nair *et al*., 2007; St Pierre *et al*., 2008). Waprins in snake venoms are hypothesised to defend the venom glands against bacterial infection (Nair *et al*., 2007) and homologs to waprins have been found in other cnidarian venoms (Klompen *et al*., 2022). Thus, these proteins may not actually contribute to envenomation itself but rather the health of the venom cells in *Acropora* and *Porites* species, making them integral to the venom defence of these species.

### Venom differences in naïve *versus* exposed corals

In this study, we included two scleractinian species that were endemic to the Caribbean and are therefore “naïve” to CoTS (i.e., *A. cervicornis* and *P. astreoides*) and nine species in Pacific reefs with a history of CoTS exposure, to examine whether venom differences may have evolved through selective pressure through predation exposure from CoTS (**Fig. 2**). Indeed, animal toxins have been shown to diversify rapidly under adaptive selection pressures (Kordiš & Gubenšek, 2000). In cnidarians specifically, the majority of toxin gene families have been shown to occur under negative selection (e.g., actinoporins and neurotoxins), where variations in venom genes accumulate rapidly under short bursts and beneficial genes get fixed in the population and accumulate for long periods of time (Jouiaei *et al*., 2015). These toxin families under negative selection are hypothesised not to be influenced by the chemical arms race that occurs between predator and prey due to their long accumulation times and high conservation of amino acid sequences (Jouiaei *et al*., 2015). One family of toxins that shows the opposite in cnidarians (and thus influenced by predator-prey interactions) is CFXs, with evidence from mixed effects models of evolution that CFXs evolved under positive Darwinian selection in cnidarians (Jouiaei *et al*., 2015). Our data would agree with these findings, where we see a large expansion of CFX homologs in CoTS-exposed corals (ranging from seven in *P. rus* to 15 in *P. compressa*), whereas *Porites astreoides* only harboured one putative CFX toxin. Furthermore, the CFX from *P. astreoides* clustered with the anthozoan JFTs, similar to *Acropora* species. Our findings of expansion and diversification of CFX homologs in the *Porites* species found across the Pacific in addition to them being related to more venomous cnidarians (box jellyfish), implies that CFXs are important toxins tailored to the geographic niche of CoTS exposure. Contrastingly to CFXs, we found *Porites astreoides* contained two conkunitzin homologs (similar to exposed *Porites* species), despite this species never coming into contact with CoTS. Therefore, these toxins may function in deterring all corallivores, rather than a specific CoTS response, agreeing with findings that neurotoxins are under negative selection pressures (Jouiaei *et al*., 2015). Thus, conkunitzins may be beneficial to all *Porites* species, regardless of environment. Research is needed to investigate these findings and compare naïve and exposed corals to understand which toxins are deployed under CoTS contact.

Apart from venom components, many other factors need to be considered to understand the complete repertoire of coral defence strategies against predation (**Table 4**). These factors include (but are not limited to) a coral species’ nematocyst community and the plasticity of this community to changing predation pressures, in addition to the metabolites and volatiles produced from coral associated taxa which may aid in deterring predators. These variations highlight the multitude of factors that must be considered when investigating the role of coral venom in defence against CoTS. Future research should aim to profile venoms and cnidomes from different polyp types of various coral species, sampled over a wide range of temperatures, locations and predation pressures, which would enable rigorous comparisons of venoms between coral species (Rachamim *et al*., 2015).

**Table 4.** Factors and variables to be considered in the future when investigating a coral’s biodefence against CoTS predation.

| Variable | Influence on a coral's biodefence |
| --- | --- |
| Number of nematocysts | Toxicity, including that inflicted from animal envenomation, is a function of exposure that is determined by both dose and duration of contact (Rozman & Doull, 2000). Therefore, corals that release more venom *per* nematocyst or have a higher density of nematocysts overall, are likely to inflict more significant damage to their predators. Nematocyst abundance differs between coral species, with *Pocillopora* species showing the highest density *per* surface area (Ben-Ari *et al.*, 2018). |
| Sensitivity of the nematocysts to mechanically discharge upon contact with CoTS | *Pocillopora* also had the most discharged nematocysts out of the scleractinian species, (although the hydrozoan *Millepora* discharging the most), whereas *Acropora*, the preferred CoTS prey, discharged almost none (Ben-Ari *et al.*, 2018). |
| Nematocyst types and diversity | Cnidarians contain between two and six nematocyst types (David *et al.*, 2008) and sea anemones toxins can be specific to certain nematocyst types e.g., Acrorhagi-specific toxins (Honma *et al.*, 2005; Macrander *et al.*, 2015). Therefore, if scleractinian species contain different types, this may alter the venom components and their sensitivity to discharge. A greater diversity of nematocyst types may allow |
|  | the coral access to an increased toxin repertoire. |
| Polyp types | Even if a cnidarian has the same nematocyst type, cnidarians such as zoanthids have been shown to harbour a different venom chemical profile, despite having the same nematocyst type (Klompen *et al.*, 2022), adding even more complexity to cnidarian venom characterisation. This difference in chemical profile could be due to differing nematocyst turnover rates or different pathways of nematogenesis between polyps (Klompen *et al.*, 2022). |
| Location of venom release from scleractinian prey | The tips of branching coral colonies have been shown to be more toxic than their base (Ben-Ari *et al.*, 2018) and as both *Acropora* and *Porites* differing in their morphology (branching *versus* massive) this may also influence the type and number of nematocysts a CoTS comes into contact with during predation, ultimately influencing venom exposure. |
| Predation pressure and previous exposure to CoTS | Nematocyst communities and abundances can vary with predation pressure e.g., *Porites compressa* colonies are able to increase their overall nematocyst density following predation from corallivorous fish (Gochfeld, 2004). |
| Novel, uncharacterised venom components | It is important to note that our study likely missed a significant portion of the venom arsenal, as we focused on known homologs from other venomous taxa and it is likely that there are novel scleractinian venom peptides and proteins that are uncharacterised and have no known homologs (Schmidt *et al.*, 2020). This underscores the importance of combining bioinformatics with experimental approaches in future work. |
| Geographic location and environmental factors | Cnidarian venoms have been shown to vary in their lethality and bioactivity within species and across geographic location and temperature (O'Hara *et al.*, 2021; Yu *et al.*, 2021). |
| Potential cytotoxic secondary metabolites and volatiles from coral associated taxa | Venom is not the only defence mechanism the coral has access to, for instance, corals and their associated communities (e.g. bacterial microbiome, fungi and dinoflagellate symbionts) contain/can synthesise a suite of volatiles (Fukatsu *et al.*, 2007; Modolon *et al.*, 2020; Chen *et al.*, 2022) |
| Symbiotic coral fauna | Symbiotic fauna (e.g., coral crabs) can also deter CoTS (Rouzé *et al.*, 2014; Deaker *et al.*, 2021). |

## Conclusion

Overall, our study provides new insights into the similarities and differences in the venom profiles between CoTS-preferred (*Acropora* species) and CoTS-non-preferred prey (*Porites* species and *E. lamellosa*) species, shedding light on the underlying biochemical factors influencing CoTS prey preferences and avoidance. Critically, our study emphasises the need for increased availability of high-quality coral genomes to help us understand genetic processes underpinning coral phenotypic traits and defenses against CoTS. While many protein families were found across all coral species, the phylogenies of these protein families and individual proteins within these families differed between coral genera. We also observed a large expansion in the diversity of these CFX proteins in CoTS-exposed *Porites* species compared to naïve *Porites* species, emphasising these proteins are likely tailored to deterring CoTS. Additionally, some proteins were found exclusively in *Porites* species venom (e.g., tereporin/conoporins, conkunitzin-S1, OVF) or *Acropora* species venom (e.g., sarafotoxin). These findings suggest that, despite similarities in the overall toxin repertoire, there are distinct molecular signatures that may influence CoTS feeding behavior. Whether these proteins contribute to a coral’s defence against CoTS predation remains to be determined and will require future research, with this study providing a foundation for such investigations. It is critical to characterise the influence of coral venom on the feeding preferences of CoTS to inform management strategies to conserve reefs during CoTS outbreaks. For instance, more venomous corals could be strategically distributed/planted throughout reefs to provide a bioactive shield for more vulnerable coral prey, inhibiting CoTS corallivory and promoting them to avail other food sources. Thus, coral venoms have the potential as an application of natural CoTS deterrents, aiding in the control of CoTS outbreaks.

## Supporting information

Publication license Figure 2

## Acknowledgements and funding sources

This research was supported by the Pacific Funds grant (HC/1674/CAB CoTS-PACIFIQUE) awarded to **SCM**, **HMP**, **LMG** and the ATER/CORAIL Laboratoire d’Excellence postdoctoral fellowship grants for “ AcantCorVenin Grant” awarded to **SCM** for **LMG**.

## Data Accessibility

All data for this study can be found at the Open Science Framework (OSF; Gorman *et al*., 2025). Previously published genomes and transcriptomes used in this study and their corresponding project accession numbers can be found in **Table 2**.

## Author Contributions

**LMG** conceptualized the experiments, designed methodology, collected and analysed the data, and constructed all figures and tables. **HMP** and **ASH** designed methodology and conducted data collection and analysis. **LMG** wrote the manuscript and **MB, SCM, HMP** and **ASH** revised the manuscript. All authors approved the final submission.

