## Supplemental File 1 for "Coral venom and toxins as protection against crown-of-thorns sea star attack"

**Table A.** Abundant toxin and peptide/protein families found in cnidarian venoms and their respective InterPro and PFAM accession numbers of their conserved domains.

| Toxin/Peptide Family | | InterPro/PFAM conserved domain and accession number | Sequence/consensus sequence used to BLAST against genomic databases (parameters: 25% identity, ignore gaps) |
| --- | --- | --- | --- |
| Pore-Forming Toxins | Actinoporins | Anemone cytotox domain - PFAM ID: [PF06369](https://www.ebi.ac.uk/interpro/entry/pfam/PF06369/) | MXRLILVIILEMDFSILCITTATALFGSPXAKXDNHVKAZXLETLLXRSXKRNSEGPEXPGSLSTNIDTEIQGRQGSSSISAXXEIKGEKRISVSKDXEKKSKSXAXAGAVITAGAXLTXXVLKRKVXVLDALGSLDNPGFTWFGLAXXXVSRKIAIGVDNPESGNYTWTAJNESVYLFRSGTSSGDNVXLPXXVRAMPXGKALLTYGARETKTXGPVATGAVGVLTYYIPDPLRLRRPLRLRIXXXXXGKTLAVMFSVPNXFDNTYNLYSNWWNVKXYXGKXRRYLADXXMYEXLYYXNAGKGXNQAXHPIFKGEXDNGWEHEKBLSPGTRYDLVCCGPSHGYSHEAGRDKPWGHMRPCKVEGYDRGEKGLKYKIYVKRGXMVYTSSGQAKLEIKVXKAXFFSQSPSKSSWWPKESETCK |
|  | *Chironex fleckeri*-like toxins (CFXs) | N/A | MXLMLXSRLFSLLLLFLFXIXJMTGIASKEXRLXRXKRSAXDTISSGLXSLKXKLDAKKPXGKQLFDXLXEXXXXXXAKPSNDDERAKVXGAXGSVGXALGKFQSGDPAKIASGCLDILXGIASTLXGPYGAKXSAVXSJLSSVXGLFSGTKAEXSVXSVVDKAFXEQRDQELQXALXGAKRXFAVSXAFLDGVRNXTSBLTPTEXSALAANVPVYQXXXFIGMXESRIXXGXPXTXLSEAKRTXXFIXLYLXLCXMRXTLLXDLIXLXXTPGGHSPNIASGIXEVSDLNKEEYXXTFEDFLHXMXXETALXGSYYYPIEHSEQSXKIFKFXXXFGVPYDDXPRXXXTGVYYRFSNRYWPNYSICKESYMGNXMFRGCSXXRYXXLRJXKLXDGYYTIKLXDGXNXYIXKHAQGWXWGTADEDPGEQGXFNFVPLKXXXRGXYMLSTKKWPNYFXYMESSASGYJRSWXXNPXKLGYGXQGHWTLXTXSNPXNXIT |
|  | Hydralysin | Anemone cytotox domain - PFAM ID: [PF06369](https://www.ebi.ac.uk/interpro/entry/pfam/PF06369/); PFM_parasporin-2-like - CDD: 380792 | KGNENKTVRXWKCAVENRSTKTLYAXGTTPDGIALNSEGYFGYHSPPITEQYGRPCYKQTGETKITSQDLSAPTDDVLGSSAAINRGDSPITLTFGVEGVFQMGGEFSLTVSVRKSGSSSVESGSMKTVLADIPPQSTGVFVWEKSRGAATGAXGVVHYXYGDKILNJMASIPYDWNLYXAWANXRVSDXKMSSEFXVEGAFKMGGEFSLTXSVGKSGSSSVEKTSTSSVQVTVPPRSKVVVSMVGIMKKEEXFSNLPVTYBGXNGAXYPTRAGNWGTVDGTKFFLTDKSHCLNKTSGVIKGTIDHANVFDVSAEFKVIFSG |
|  | Membrane-attack complex/perforin (MACPF) toxins | [PF01823](https://www.ebi.ac.uk/interpro/entry/pfam/PF01823/) | ASSEEEVQGLSLNLKAYSMSSILKKNCVNTKPLSKDLVSDFEALDSEITKPWKLSSWKKYKVLLEKYGSHIVKESISGSSIYQYVFAKSSQKFNHRSFTVKACVSLAGPTXVGKLGFSGCTGVSQQEIEQSSSQSMIKKLVVRGGKTETRASLIGELDPDQINKFLIEAETDPSPIQYKFEPIWTILKXRYVGTEHFAKAVNL |
| Enzymes | Phospholipase A2 | PF00068 | LLQFXXMIACXTGRLSXXDYXGYGCWCGLGGKGTPVDDVDRCCYVHDXCYNXIXXGPRPTCSPDKXAXYXKNYXTSKRRGCLKCSXXSXXXCLXWXTSKCXRXJCXCDIXAAKCFARNHFNPLQDKYK |
|  | Zinc Metalloprotease (conserved zinc binding site highlighted in orange) | Peptidase M10 - PFAM: PF00413; Hemopexin - PFAM: PF00045 | ADFSPSDSVKRKRRYXXXGMYTKXLWNKXXITWXVXNDNNDGISXEXVAXXXXERALXKWSXVTXJRFXERKNLXXVNAXXPDIXXRFGAWRETVLFXRKXHGDXQLMTAYPFDGXGGXVLAHAFYPXXSXTPLXGDVHXDDDXXEVFXIXSNXSGSGKDLFILLWXVVHELGHSJGLDHSDXRXXXMYXIXSPXYRGXXGLDFXLTXDDIXGAQSLYGXKXXPTXKXPXPXPGNRFDKXXKPXDAXCXXKXGAXXLIKEGNXKRTYVFNXDKXYILNXXDLXVDXGPIXVSSXFXGXXXVDAAFXRQXDXXXXXFSGNSYYVXSXGXDYDXVXGPXXISDGFXGLPANFGDIDAAFCKVWPGNNXXYIFKGXDYWRFYQXXNQNAYQADSGYPRKIRXAWIGXPDXXDAVXXWXNGXXYFFKGSKYYRINKNRDMDDYCFNVEXGYPRXXTEXXLYCXS |
| Neurotoxins IPR000693 | ATX-III domain | PFAM ID: PF08098 | AGGKXXCCPCAMCKYXXGCPWGQXXXXXGCSXPKV |
|  | Defensin domain | Superfamily database ID:SSF57392; PFAM ID: PF07936 | KRGVPCLCDSDGPXVRGNTLSGTVWLXTGGYGXGGCPSGWHKCKSYFGPXIGWCC |
|  | Potassium type 5 neurotoxin | None | EKRCKGKYQKCTSNSDCCDXKDRAGRRKLRCLTQCDEGGCLKYKQCLFYXGLQ |
|  | Potassium type 6 neurotoxin | None | RCKTCSKGRCRPKPNCG |
|  | Kunitz-type domain | PFAM ID: PF00014 | INKDCLLPXDVGRCRARFPRYYYNSSSRRCEKFIYGGCGGNANNFHTLEECEKVC |
|  | Shk domain | PFAM ID: PF01549 | CKDNFPXSTCXHVKXNGNCRTSQKYRXXNCAKTCGXC |
|  | Small cysteine rich peptides (SCRiPS) | None | MGVKFNLCLLLLLXXXXSXQGXXXXKKXDSTDEKXFHGNYRRXXXCDXPXGVCYYIHDRCPPGTPXDCGQXXFGCYLPTNRCCC |
|  | Turripeptide from *Millepora alcicornis* | Kazal_2 domain (PFAM ID: PF07648) | MSILTLSVTIALFGNTASFITRKCQKACPLIYNPVCGSDGVTYPNQCALDVATCESNGKITKVSDGPCSKCQKACPLIYNPVCGSDGVTYGNQCALDVATCKSDGKITKVSDGPCSTL |

**Phylogenetics**

**Actinoporin tree**

Actinoporins discovered in the genomes of the *Acropora* and *Porites* species and the transcriptome of *E. lamellosa* (**Table S1**) were aligned with previously discovered scleractinian (Ben-Ari *et al.*, 2018) and anthozoan (Frazão *et al.*, 2012) actinoporins, in addition to related sister cytolysin proteins from the same superfamily (e.g., hydralysins; Sher *et al.,* 2005a, b) and tereporins (Gorson *et al.*, 2015), using MUSCLE in Geneious Prime (version 2024.0.7). Alignments were then manually edited and trimmed to the anemone cytotox domain (PFAM ID: PF06369), resulting in an alignment of 94 sequences and 582 aa (amino acids) in length. The alignment was submitted to IQ-tree (Trifinopoulos *et al.*, 2016) to obtain the correct model of amino acid evolution. Using the corrected Akaike information criterion (AICc), the best-fit model of evolution was WAG+G4 with a log-likelihood score of -15500.97. Once this model was obtained, maximum likelihood trees were generated in IQ-tree using 1000 ultrafast bootstrap replicates and represented as SH-aLRT support (%). The actinoporin tree was rooted with the plant bryoporin Physcomitrin from *Physcomitrium patens* (AAV65396), which is related to actinoporins (Šolinc *et al.*, 2022). Trees were edited and visualised in Interactive Tree of Life (iTOL; version 6; Letunic & Bork, 2024).

**MAC-PF tree**

We constructed a MAC-PF tree separately from the actinoporin tree due to the large number of putative MAC-PF’s found in this study. However, we did include several confirmed anemone actinoporins in the MAC-PF tree as MAC-PF’s and actinoporins are distantly related and share structural similarity, forming distinct sister clades to one another (Frazão *et al.*, 2012). Thus, we included a subset of actinoporins from Frazão *et al.* (2012) to create the correct tree topology: RTX-A from *Heteractis crispa* (P58691); Tenebrosin C from *Actinia tenebrosa* (P30833); and PSTX-20a from *Phyllodiscus semoni* (P0DL55). Alignments were generated using MUSCLE in Geneious Prime (version 2024.0.7). Alignments were then manually edited and trimmed to the MACPF domain (PFAM ID: PF01823), resulting in an alignment of 72 sequences and 269 aa in length. We removed nine of the *E. lamellosa* putative MAC-PF sequences as they did not contain the MAC-PF domain (PFAM ID: PF01823; **Table S3**) and they aligned with confirmed MAC-PF’s downstream from the MAC-PF domain, emphasising that they are likely partial sequences. Furthermore, these putative partial *E. lamellosa* MAC-PF’s did not group with any other MAC-PF’s or actinoporins in the MAC-PF tree, disrupting the tree topology. The final alignment was submitted to IQ-tree (Trifinopoulos *et al.*, 2016) to obtain the correct model of amino acid evolution. Using the corrected Akaike information criterion (AICc), the best-fit model of evolution was WAG+G4 with a log-likelihood score of -10208.8217. Once this model was obtained, maximum likelihood trees were generated in IQ-tree using 1000 ultrafast bootstrap replicates and represented as SH-aLRT support (%). We rooted the tree with the same protein as the actinoporin tree - plant bryoporin from *Physcomitrium patens* (AAV65396). Trees were edited and visualised in Interactive Tree of Life (iTOL; version 6; Letunic & Bork, 2024).

**CFX tree**

CFXs retrieved from custom BLAST and discovery bioinformatics were aligned using MUSCLE with 71 jellyfish toxins (JFTs) and their homologs, including CFXs, retrieved from Klompen *et al.* (2021). Alignments were then manually edited and trimmed. As the CFXs had no conserved domains, the alignment was trimmed to an area showing the most conserved residues across species, resulting in a consensus alignment of 118 sequences and 594 aa in length. The alignment was submitted to IQ-tree (Trifinopoulos *et al.*, 2016) to obtain the correct model of amino acid evolution. Using the corrected Akaike information criterion (AICc), the best-fit model of evolution was LG+F+G4 with a log-likelihood score of -28977.45. Once this model was obtained, maximum likelihood trees were generated in IQ-tree using 1000 ultrafast bootstrap replicates and represented as SH-aLRT support (%). We rooted the tree using the bacterial cry-like toxin Cry2Aa from *Bacillus thuringiensis* subsp. *kurstaki* (P0A377) to root the CFX tree as CFX’s have structural homology to insecticidal Cry-like toxins (Brinkman *et al.*, 2014; Klompen *et al.,* 2021).

**SCRiPS tree**

A SCRiPS tree was constructed based on recent data published by Barroso *et al.*, (2024), dividing scleractinian SCRiPS into four phylogenetically distinct clades (*SCRiP-α, SCRiP-β, SCRiP-ϒ and SCRiP-δ*). We extracted several protein candidates for each SCRiP clade from Barroso *et al.*, (2024) to align with our putative SCRiPS hits resulting in a consensus alignment of 128 sequences and 156 aa in length. The alignment was submitted to IQ-tree (Trifinopoulos *et al.*, 2016) to obtain the correct model of amino acid evolution. Using the corrected Akaike information criterion (AICc), the best-fit model of evolution was PMB with a log-likelihood score of -7416.135. Once this model was obtained, maximum likelihood trees were generated in IQ-tree using 1000 ultrafast bootstrap replicates and represented as SH-aLRT support (%). The tree was rooted with a SCRiP from the octocoral *Scleronephthya gracillima* (GJHC01022697.1.p2; Barroso *et al.*, 2024).
